# Large-scale ciliary reversal mediates capture of individual algal prey by Müller’s larva

**DOI:** 10.1101/709790

**Authors:** George von Dassow, Christina I. Ellison

## Abstract

We documented capture of microalgal prey by several species of wild-caught Müller’s larvae of polyclad flatworm. To our knowledge, this is the first direct observation of feeding mechanism in this classical larval type. High-speed video recordings show that virtually all captures are mediated by large-scale transient ciliary reversal over one or more portions of the main ciliary band corresponding to individual lobes or tentacles. Local ciliary beat reversals alter near-field flow to suck parcels of food-containing water mouthward. Many capture episodes entail sufficient coordinated flow disruption that these compact-bodied larvae tumble dramatically. Similar behaviors were recorded in at least four distinct species, one of which corresponds to the ascidian-eating polyclad *Pseudoceros*.

## 1. Introduction

Müller’s larva of certain polyclad flatworms is one of the iconic invertebrate larval forms, yet its way of life in the plankton remains poorly known. Lacking internal cavities other than the stomach, its bilaterally-symmetrical body is elaborated into several heavily-ciliated lobes and solid arms. It has eyes and brain, apical and caudal cirri, and swims rapidly with a rifling motion. (A similar, less-elaborate form known as Götte’s larva, is herein treated as a simpler but functionally equivalent form, or as a developmental stage.) Like many larvae that feed on phytoplankton, Müller’s larva possesses a convoluted band of denser ciliation that loops around the mouth (Lacalli, 1982). Like many ciliated invertebrate larvae, it is inferred to feed in the plankton because it appears in a spectrum of sizes from which growth is inferred, and has been found with apparent food in its stomach (reviewed in Nielsen, 2005). Yet only recently was any Müller’s larva definitively shown to feed and grow on unicellular algae (Rawlinson, 2010; Allen et al., 2017). How they do so remained unknown until now: based on high-speed video recording of individuals from natural plankton, we show here that Müller’s larvae of several species capture individual algal cells, one by one, using a large-scale transient ciliary reversal coupled to muscular flexions that together bring food-containing fluid parcels toward the mouth – or rather, as will be seen, bring the mouth to the food.

## 2. Materials and Methods

### 2.1 Organisms and culture

Wild Müller’s larvae were collected in the Charleston, Oregon marina by dockside plankton tow using a 53 μm net. These larvae are positively phototactic, and can therefore be concentrated by selecting the fraction that collects in the jar at the window-side surface, then isolated by pipette into filtered seawater in custard bowls placed in seatables. Larvae were supplied with >10^4 cells/ml *Rhodomonas* cf. *lens* (CCMP739) or *abbreviata* (CCMP1178), sometimes supplemented with *Prorocentrum “micans”* (obtained from Carolina Biological) or *Pyramimonas parkae* (CCMP725) or other microalgae. All algal cultures were maintained in L1 medium made in microfiltered natural seawater, and were *not* centrifuged before feeding experiments, so as not to deplete ejectisomes. Larvae could be maintained for two weeks or so on *Rhodomonas cf. lens* or *abbreviata* (which is larger and better armed with ejectisomes) and were observed to grow somewhat and add more lobes. Some underwent an irreversible metamorphosis to swimming and crawling worms. As described previously for *Stylochus elliptica* (Allen et al. 2017), only larvae fed on cryptophytes grew, although they clearly ingested some other algal cells. For comparison to wild-caught larvae, egg plates laid by *Pseudoceros canadensis* were peeled off colonies of *Distaplia occidentalis* (also obtained from the Charleston marina) and allowed to hatch in bowls of filtered seawater.

### 2.2 Video recording

For recording feeding behavior, larvae were placed in cuvettes made from slide and coverslip separated by various shims (clay feet, slide fragments) thick enough to allow them to tumble freely. Alternatively, we placed them in coverslip-bottomed dishes with a coverslip on top of the well. To prevent evaporation, chambers were sealed by painting around the edge with molten vaseline. For some experiments, the seawater was seeded with a dilute suspension (~1:10^4) of half-and-half (Umpqua Dairy) to provide a neutral tracer of fluid motion. For tethered preps, larvae were caught by capillary suction on the end of a fire-polished micropipette with a ~50 μm aperture made from 1.0 mm x 0.5 mm microinjection capillaries using a Sutter P1000 horizontal pipette puller and a Narishige MF-9 microforge.

All recordings were made on an inverted microscope (Leica DMi8) using DIC optics and a high-speed camera (Photron Mini UX100) recording at 500 or 1000 frames per second with 1 or 0.5 ms exposure. This was sufficient to stop most ciliary motion. Note that due to the folded light path of the inverted microscope, all images are mirror images; to save processing steps we have not reversed them. Although we have detected no handedness to behaviors reported here, it is likely that since at least the rotation of larvae as they swim has a handedness, so may prey capture functions. Any future use of our video data may need to take this into account.

### 2.3 Image processing

Image processing was performed using ImageJ, and figures were composited in Adobe Photoshop CS6. In some figures, as noted in legends, Fourier filtering was applied to isolate high-frequency image elements (i.e., particles). No other convolution filtering (sharpening, noise reduction, etc.) was applied to any image. Other operations included stack registration to change from laboratory to larval reference frame, reslicing to create kymographs, and frame averaging or projection to visualize pathlines. All kymographs were created by automatically registering video frames (using the StackReg plugin to ImageJ) to convert the sequence to a larval reference frame, fitting a spline to the profile of a ciliated zone, applying the Straighten function within ImageJ, then replacing the result. In most kymographs, to eliminate confusing shadows and glows from the larval body or passing cells, the resliced data were high-pass filtered using the FFT functions within ImageJ with a band from 0 to 4 pixels.

### 2.4 DNA barcoding

Individual larvae were photographed, rinsed several times in micro-filtered seawater, and frozen in 1-2 μl seawater at -80 C for DNA extraction. DNA was extracted from adult tissue with DNEasy Blood and Tissue kit (Qiagen), and from larval tissue with Chelex matrix (InstaGene, BioRad). DNA from adult *Pseudoceros* required 100-fold dilution to allow PCR. Barcoding regions of two mitochondrial genes, 16S rRNA (448-bp) and COI (658-bp) were amplified using universal primers LCO1490/HCO2198 (Folmer et al., 1994), 16SARL/16SBRH (Palumbi et al., 1991) and universal variants jgLCO1490/jgHCO2198 (Geller et al., 2013). PCR products were confirmed using gel electrophoresis, purified using Wizard SV Gel and PCR Cleanup kit (Promega), and sequenced directly (Sequetech) using both forward and reverse primers. Sequences were analyzed using Geneious version 2019.0 (Biomatters). Forward and reverse sequences were trimmed, aligned, and proofread. Resulting consensus sequences were aligned (MAFFT alignment), and used to build neighbor joining trees. Alignments and trees were built using default parameters. COI sequences were translated to amino acid sequences within Geneious to check for stop codons. Only sequences with no stop codons in the reading frame were used in the analysis. Sequences have been deposited in Genbank (accession numbers in Supplemental Table 1)

The vast majority of video recordings were made with the medium-sized, wild-caught larvae of unknown parentage (what we call the Winnie-the-Pooh morphotype), because their relatively simple form when young, modest swimming speed, and eagerness to feed made them good subjects. Nevertheless we observed similar behaviors by eye, free-swimming in bowls, for all morphotypes and for hatched *Pseudoceros*, and obtained high-speed recordings of successful captures from all but one morphotype (which we refer to as Broad-bill) which did not perform under bright lights while trapped in cuvettes.

## 3. Results and Discussion

### 3.1 Observing prey capture

When trapped under a coverslip, Müller’s larvae transform into a bullet-shaped fast-swimming posture, and may remain in this form for many minutes. Even when provided with food in tightly coverslipped preps, they were not observed to feed or to react in any way to passing prey. Yet when observed unconstrained in a dish and provided with unicellular algae *(Rhodomonas* or *Prorocentrum*), they relaxed into a canonical lobate form, and although they swam rapidly, they exhibited frequent tumbles and subtler direction changes that appeared to be related to feeding. When provided with a high density of heavily-pigmented prey cells *(Rhodomonas*), our Müller’s larvae clearly ate them, as they accumulated in their guts the characteristic magenta of a *Rhodomonas* diet.

To observe the mechanism of feeding, we placed one or a few larvae in cuvettes with ample prey and enough space that larvae could tumble freely. Although incredibly inconvenient to the microscope operator, this extra space is essential: even slight constraint made it impossible for these larvae to capture prey, for reasons that will become obvious from the account to follow. Unconstrained larvae generally relaxed within 5-15 minutes, and, with sufficient suspended prey, began to feed. To record prey capture, we followed larvae around the cuvette or dish using the stage control. We detected capture events on the real-time feed from the high-speed camera by changes in trajectory. Indeed, upon playback, most abrupt changes in direction proved to involve attempted prey capture. Essentially all prey capture events we recorded were accompanied by a) transient reversal of ciliary beat in one or more lobate zones and b) lateral flexure of the oral lobes to widen the slot around the mouth (Fig. 1; Videos 1, 2, and 3). Observed ciliary reversal episodes lasted on the order of hundreds of milliseconds (i.e., several beat cycles) and involved on the order of hundreds of microns of ciliated band (i.e., most or all of an individual lobe). Because many captures involve only subtle direction changes, especially for larger larvae with more complex shapes, it is likely that we missed many of the simpler events that involved more transient reversals. *Our description herein relies entirely upon video sequences of larvae, their prey, and cilia in motion; these data translate poorly to two-dimensional representations, so we urge readers to use our figures as interpretive guides to the videos published along with this paper*.

**Figure 1.**
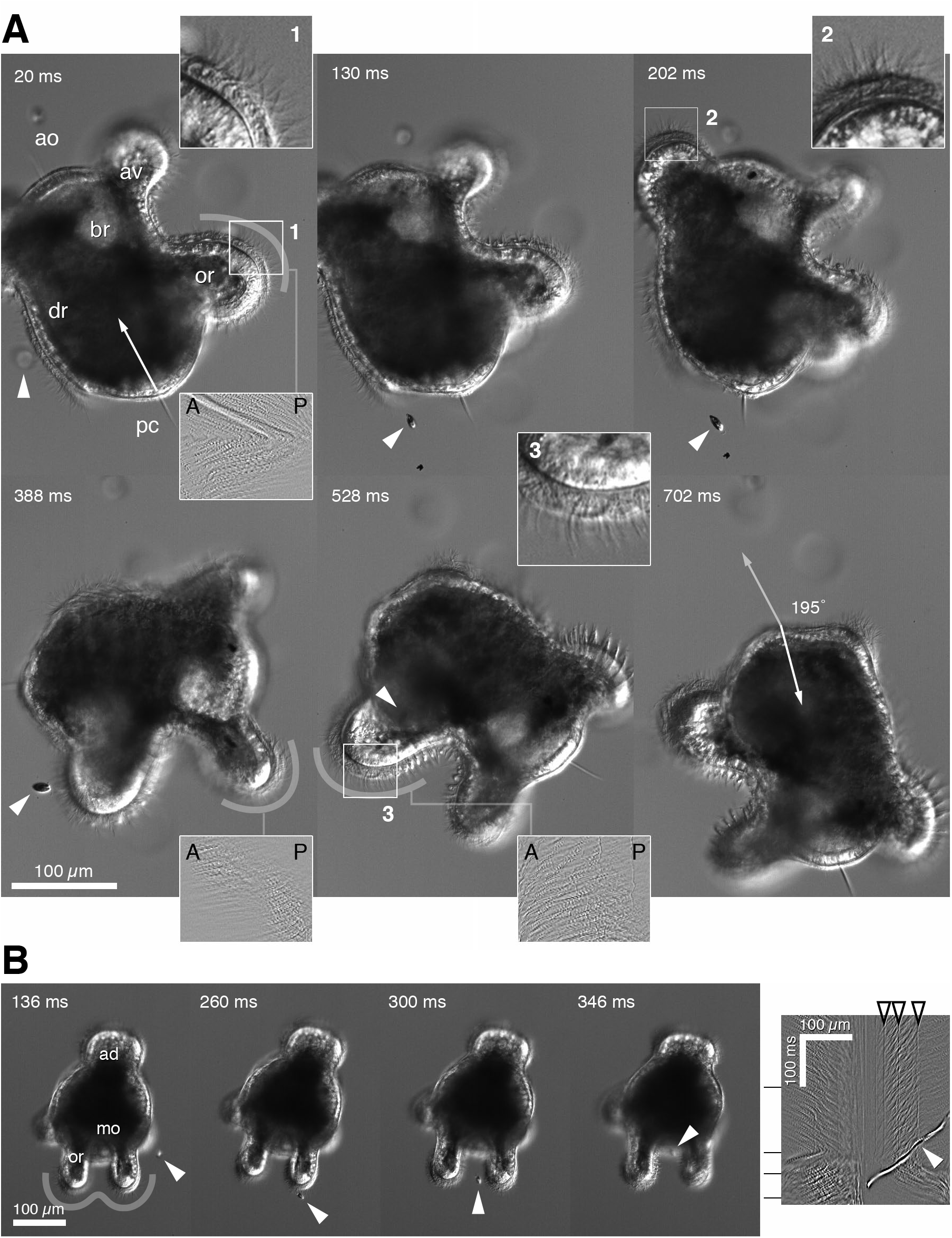
Capture of microalgal prey by Müller’s larva. A) Frames from a loosely-coverslipped capture sequence filmed at 500 fps. The captured cell is indicated by a white arrowhead: this cell arrives near the dorsal posterior of the larval body, then the larva turns until the prey is aligned with the slot between the oral lobes. Arrows indicate the initial and final swimming orientation. Insets provide 2.5x-magnified views of 1) oral lobe before capture – curved recovery profiles indicate posteriorward beat; 2) anterodorsal lobe after capture maneuver begins – continued posteriorward beat; 3) oral lobe during capture maneuver – reversed ciliary beat. Kymographic insets are made from the approximate strips as indicated over 200 ms interval, high-pass filtered to isolate ciliary strokes. All kymographs are oriented the same, such that a left-to-right diagonal indicates normal posteriorward beat, whereas right-to-left diagonal indicates reversed beat. The beat reversal in shown in the first frame is abrupt, and continues for several hundred ms (through the fifth frame); the anteroventral lobe reverses with the oral lobes in this event. B) Lower-magnification view of a capture sequence filmed in a dish, with nearly straight-on posterior view. In this case, the prey cell encounters the oral lobes first. Kymograph is made from the gray strip shown in the first frame. In the kymograph, the bright trace is the path of the captured cell (white arrowhead in frames also). Open arrowheads are the traces of stationary cilia. In this view, oral lobes beat toward the midline, until shortly after prey encounters stationary cilia, whereafter both lobes reverse their beat. ao=apical organ, ad=anteordorsal lobe, av=anteroventral lobe, br=brain, dr=dorsal ridge, mo=mouth, or=oral lobe, pc=posterior cirrus.

### 3.2 Flow around Muller’s larva

Deducing the mechanism of particle capture requires a map of flows driven by the ciliated band around the larval body. While swimming in canonical posture, Müller’s larva extends an anterodorsal and anteroventral lobe (the latter like an awning above the mouth) and at least one pair of oral lobes that are held together with a narrow slot between them, deep within which lies the mouth (Fig. 1B; diagrammed in Fig. 2A,B). All of these lobes are muscular and can be flattened, flexed, or withdrawn, and along their outlines is stretched a broad band of dense ciliation. As larvae grow, more lobes elaborate from the posterior portion of the dorsal loops of ciliated band. Some forms elongate lobes into tentacles, or elaborate anterior lobes into multiple flaps.

**Figure 2.**
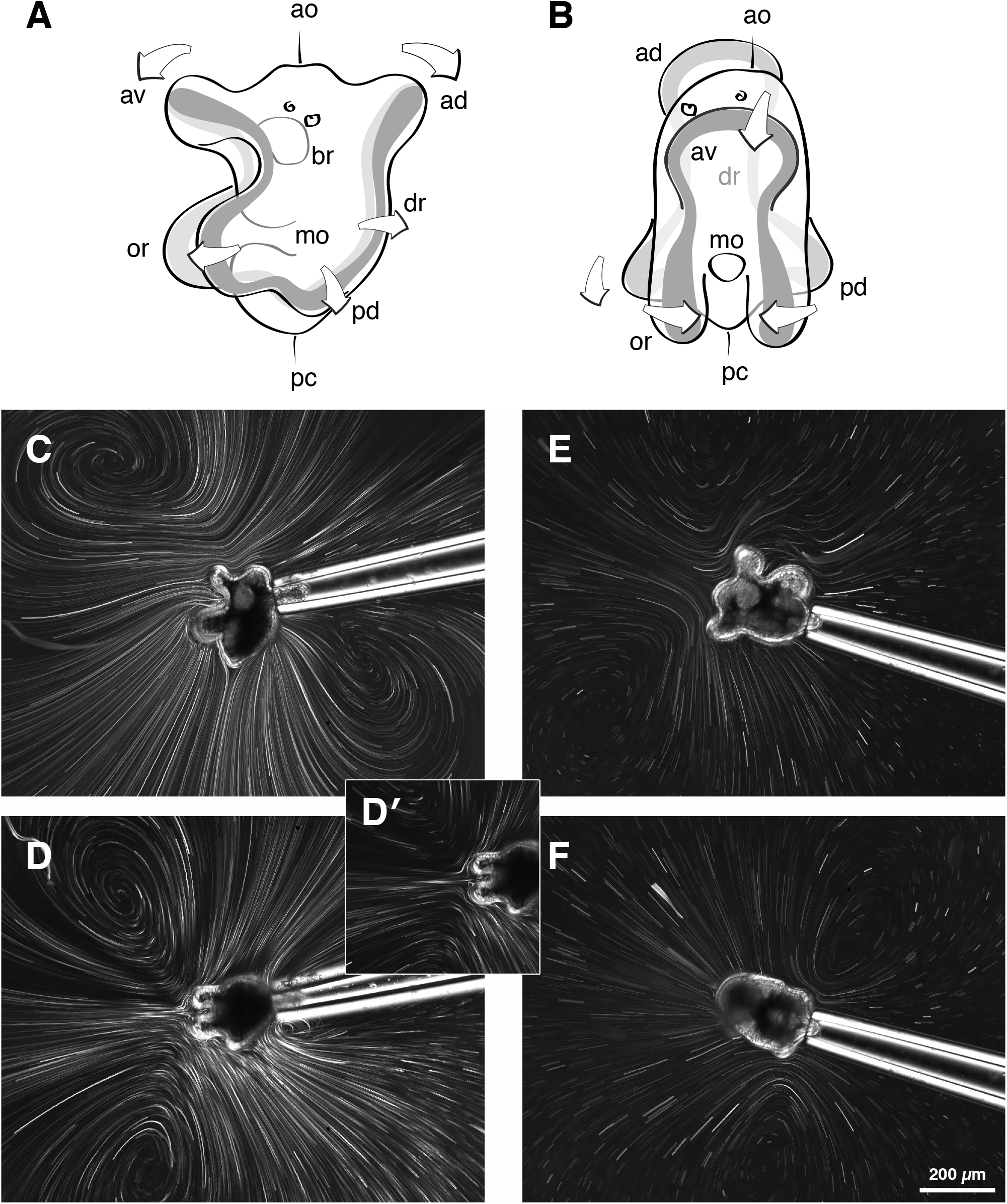
Flow fields around Müller’s larva. A,B) drawings depicting the course of the ciliated band in relation to mouth and lobes, in side and frontal view. Note, only two eyes are depicted, but many of the larvae we used had three (or possibly more; they are difficult to count in highly-pigmented individuals). ao=apical organ, ad=anteordorsal lobe, av=anteroventral lobe, br=brain, dr=dorsal ridge, mo=mouth, or=oral lobe, pc=posterior cirrus, pd=posterodorsal lobe. C-F) Flow fields around larvae tethered by capillary suction and suspended >500 μm from boundaries in seawater seeded with half-and-half. C (side view) and D (posterior view), both held by anterodorsal lobe, cover 4 s of time during which no interruptions or reversals occurred; D’ is a 400-ms subset to show individual pathlines around the oral lobes. E (side view) and F (dorsal view) are 400-ms projections from the same larva held by the posterior end (hence rotating on the tether). These emphasize pathlines around anterior lobes and posterodorsal lobes. Vortices are artifacts of nearby boundaries, but near-field flow directions are faithful.

To visualize flow we used gentle suction to tether larvae to fire-polished capillaries, in cuvettes deep enough to allow more than a body diameter to the nearest wall (~0.5-1 mm). We seeded the medium with a dilute suspension of half-and-half, then projected recordings over various time intervals to describe pathlines around tethered larvae. Cilia on both anterior lobes beat posteriorly, but with a dorsal or ventral component that varies with posture (Fig. 2C,E). Cilia on the oral lobes beat diagonally, with posterior and medial components, such that a current emerges ventrally from the colliding streams driven by the lobes (Fig. 2D). Cilia along the dorsal ridges beat toward the dorsal midline, with only a slight posterior component, and the nascent posterodorsal lobes beat directly posteriorly (Fig. 2F). Pathlines around tethered animals are compressed in the axis of flow compared to what would be observed around freely-swimming larvae (Emlet, 1990), and artifactual vortices due to nearby boundaries are unavoidable (Pepper et al., 2009; von Dassow et al., 2017). Nevertheless, this method reveals the near-field flows. In exceptional cases we managed to catch short sequences of untethered larvae swimming nearly straight, far from boundaries, and in focus for long enough to confirm the profile derived from tethering (Supplemental Fig. 1).

### 3.3 Ciliary reversal

The local ciliary reversals accompanying prey capture follow a common pattern depicted in Fig. 3A: the steady forward swimming beat appears to involve a small fraction of the cilia at any instant; reversal is sudden across a large portion of the visible lobe, and involves a much larger fraction of the cilia; thereafter, restoration of the normal beat involves a large fraction of visible cilia, including zones that prior to reversal seemed almost completely inactive. Although we did not succeed to observe successful prey capture by tethered larvae, we did record instances of reversal, as shown in Fig. 3B, which showed that beat reversal suffices to elicit large-scale flow reversal near the altered zone. On untethered larvae, recordings in the presence of neutral fluid tracers demonstrated that reversal accounts for the orientation changes that accompany capture (Fig. 4A and Video 4): in such sequences, when images are aligned from laboratory to larval reference frame, the reversal causes the larva to orbit within the local fluid domain. Perhaps this seems unsurprising – what else could account for direction change? – yet we note that this consequence would *not* be readily observed in other larvae, such as echinoderm dipleurulas, that feed using ciliary beat reversal. This is likely because their large bodies require much more impulse to steer, whereas Müller’s larva is compact. In effect, Müller’s larva orbits itself around a parcel of food-containing water. Revisiting the video accompanying Fig. 1 (and those summarized in Fig. 5) emphasizes this point.

**Figure 3.**
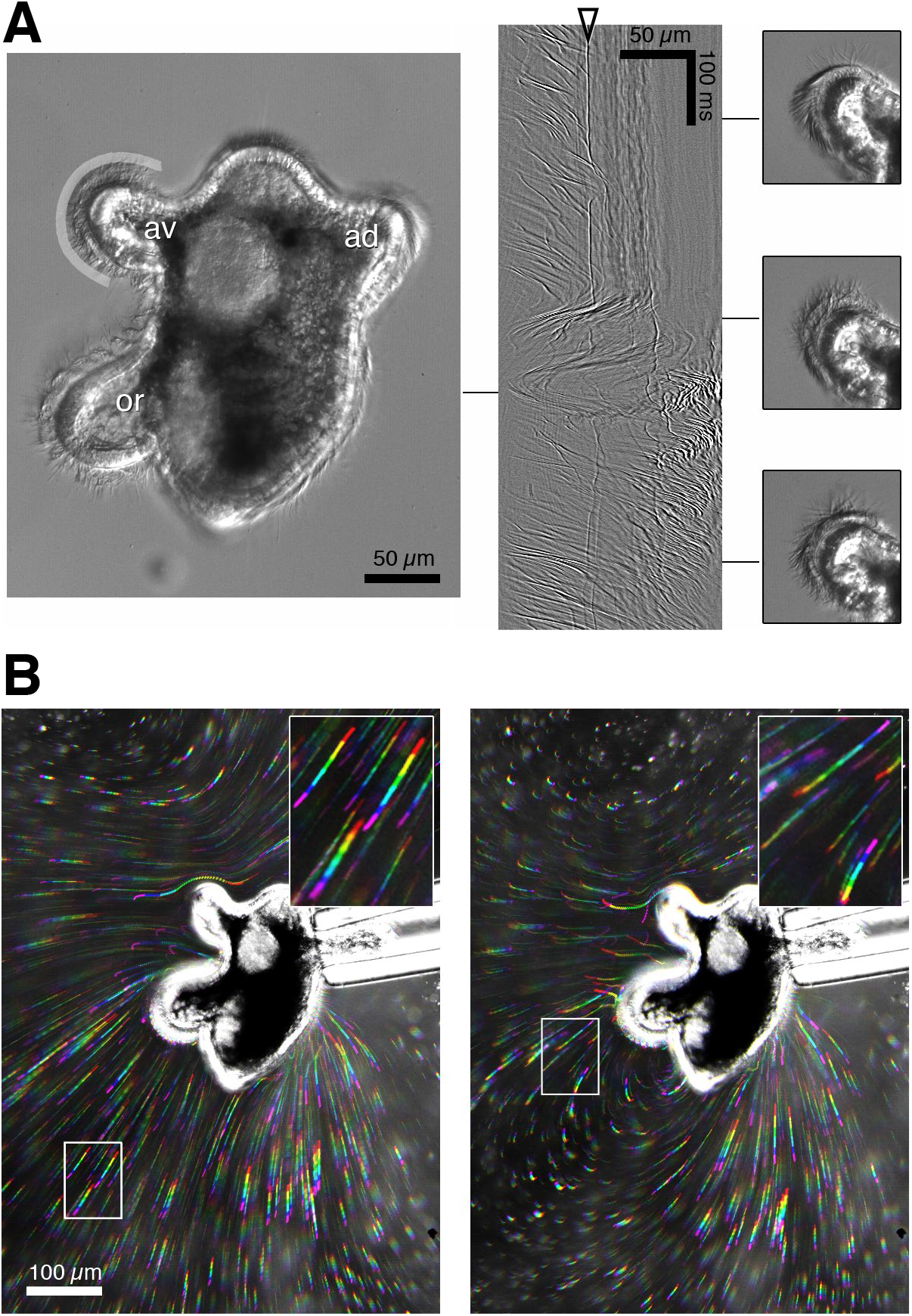
Ciliary reversal. A) Side view of an individual that exhibits a stereotypical reversal and restoration without changing focus. Kymograph is made from the profile traced in gray around the anteroventral lobe; insets of the lobe itself correspond to times in the kymograph as indicated, and show the following traits: before reversal, few cilia beat at any instant, and most appear in recovery posture; during reversal, far more cilia beat, and from this perspective appear to do so almost synchronously in a given transverse focal plane (hence kymograph traces show two distinct sweeps by many individual cilia) – note that in the orthogonal focal plane, this would appear as a strong metachronal wave *along* the band; after restoration of normal beat direction, more cilia tend to beat at any instant than before reversal. Open arrowhead points out the trace of a stationary cilium. ad=anterodorsal lobe, av=anteroventral lobe, or=oral lobe. B) Side view of pathlines during normal beat (left) and during joint reversal of anteroventral and oral lobes (right), in a tethered larva (as in Fig. 2). Rainbows are 120-ms projections, with successive 20 ms segments colored such that red is the earliest and magenta the most recent segment. Reversal draws fluid from a large fan-shaped sector toward the mouth and, in this case, over the anteroventral lobe; dorsal ridges and posterodorsal lobes maintained normal beat here.

**Figure 4.**
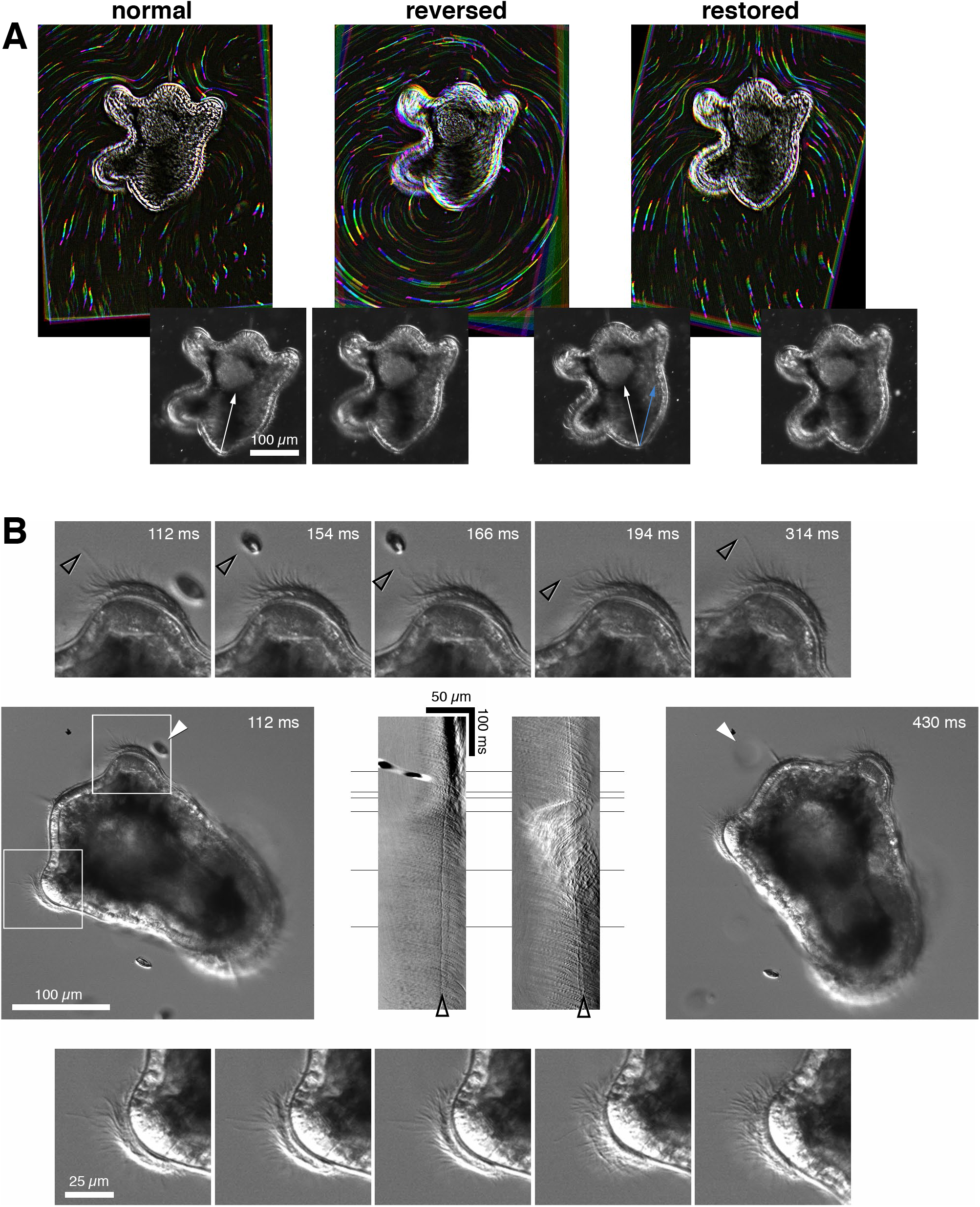
Ciliary reversal orbits the larval body within its medium. A) A transient reversal as a “free-swimming” larva catches a cell by the anterior-ventral route. This larva is more than one body diameter from any boundary, which we know because no out-of-focus stationary particles are present in the original recording. Frames at the bottom show laboratory reference frame; this larva executes a modest turn. The frames at the top have been first registered to shift to the larva’s reference frame, then high-pass filtered to highlight small particles, then rainbow-coded as in Fig. 3B. B) A rare view of a very loosely-coverslipped larva in nearly perfect dorsal view. In this sequence, the prey first encounters the left posterodorsal lobe. Boxed regions are displayed at 2x magnification in sequences above and below; kymographs (without high-pass filtering) from left and right lobes in between the initial and final frames – in both kymographs, anterior is left and posterior is right, as in Fig. 1A, hence left-to-right diagonals indicate normal beat, right-to-left diagonals indicate reversed strokes. Hollow arrowheads point out traces of stationary cilia. Insets and kymographs show that whereas the prey cell encounters and deflects a non-beating cilium on the left lobe, the left lobe maintains normal beat; reversal takes place on the right lobe within 10 ms of the prey encountering the non-beating cilium. The white arrowhead indicates the prey cell before encounter, then after turning (now out of focus) aligned with the oral approach zone.

**Figure 5.**
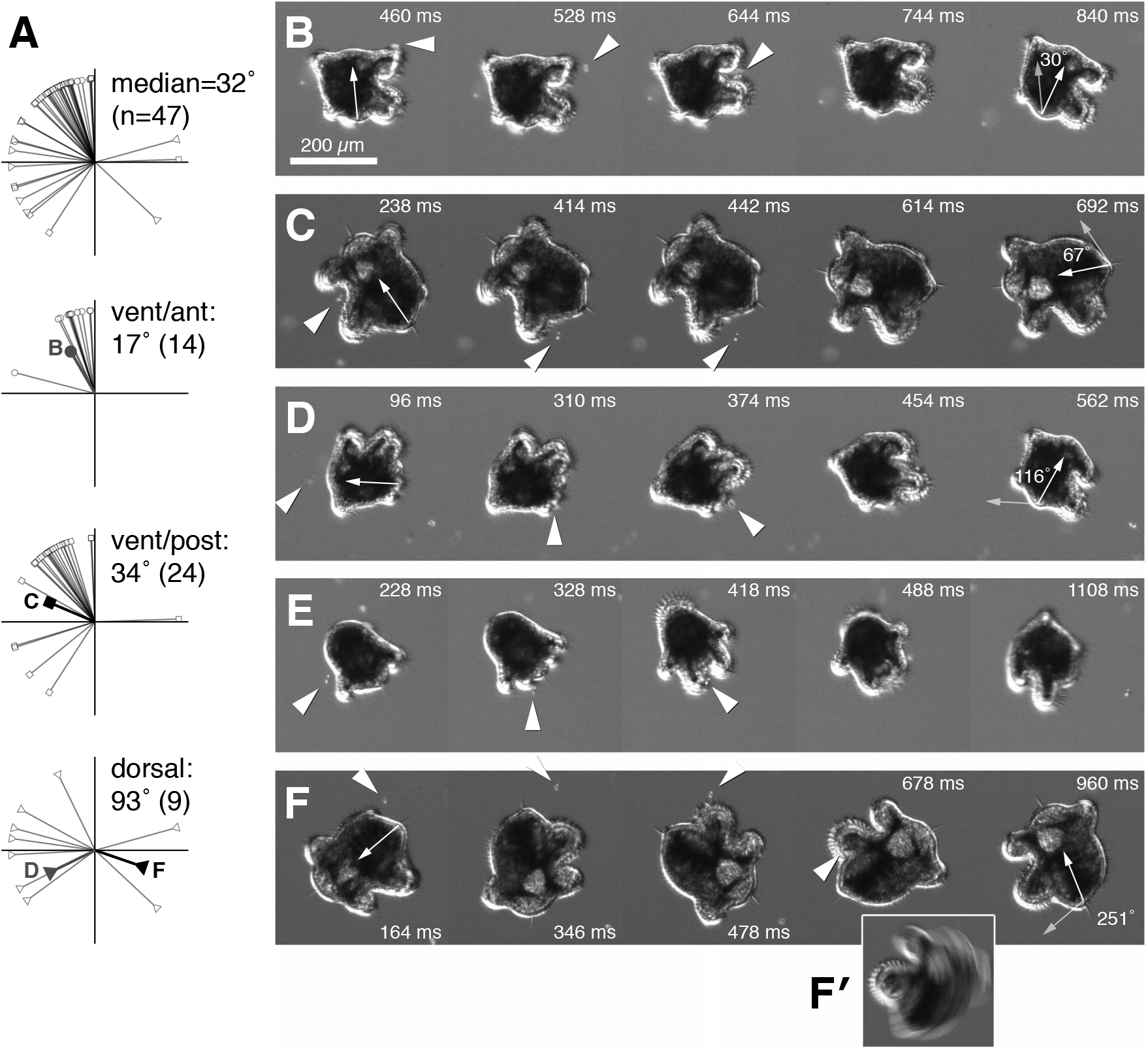
Prey capture is accompanied by turns and tumbles. A) Plots show the angle of turn measured from a common set of 47 recordings, all at 20x and featuring the Winnie-the-Pooh morphotype, in which the larva remained roughly in the focal plane throughout the event. Each event is categorized by route: circles for the ventral-anterior route (most straightforward), squares for the ventral-posterior route, and triangles for the dorsal-posterior route (which often involves dramatic tumbles). B–F) Low-mag sequences of capture events far from boundaries (as shown by the absence of stationary out-of-focus particles). B, D, and E feature individuals of the *Pseudoceros* morphotype; C and F feature Winnie-the-Pooh type. B) the ventral-anterior route; C) ventral-posterior route; D) dorsal-posterior route; E) dorsal-posterior route in initially dorsal view; F) dorsal-posterior route. For comparison to the data from coverslipped larvae in (A), the sequences in B, C, D, and F are overlaid on appropriate plots in panel A as well; note that these few free-living turns each exceed the median for constrained turns of similar category. F’) A 30-frame (= 60 ms) average from mid-turn: the oral lobe and its cilia are in nearly sharp focus, indicating that they are stationary relative to the microscope as the larva pivots.

What prompts reversal? Clearly, reversal follows the approach of prey to the ciliary band. In all the Müller’s larvae we have observed, single non-beating cilia are dispersed at regular intervals amongst the much greater number of beating ones. These are collar cells (Lacalli, 1982; Semmler and Wanninger, 2010), reminiscent of the organization described for the nemertean pilidium larva (von Dassow et al., 2013; Hindinger et al., 2013) and for actinotroch (Hay-Schmidt, 1989) and brachiopod (Strathmann, 2005) tentacles (among others), and in those other animals the collar cells are presumed to be sensory. In some video sequences of Müller’s larvae, the prey item encountered the ciliated band in focus, and could be seen to physically contact a non-beating cilium and deform it transiently (Fig. 4B and Video 5). Ciliary reversal followed within milliseconds. Remarkably, in the sequence depicted in Fig. 4B, the reversal takes place *not* in the lobe where the prey was first encountered, but *on the opposite side of the larva body*, ~100 μm away. Why? In this particular case, the reversal on the far side, with continued normal beat on the detecting side, causes that larva to orbit such that the prey item is nearly aligned with the slot between oral lobes (not directly visible in this sequence, but position is inferred); there must follow reversal of oral lobe beat to draw food-containing water toward the mouth. Such choreography bespeaks an organism wired for control of local flow with millisecond and micrometer precision.

### 3.4 Prey capture routes

We noticed that prey traversed several routes, and that these correlated with the extent of ciliary reversal. Some prey cells approached along paths from the anterior ventrally; these were captured by a brief reversal of ciliary beat on the oral lobes, outward flexure of the lobes, and the least change in swimming direction on average (Fig. 5B,C and Video 6; measured direction change in Fig. 5A, plot 2). In some observed instances, cilia reversed on just the oral lobes; in others, ciliary beat reversal took place on both anterodorsal and anteroventral lobes. The latter cases seemed to result in a transient stall of larval swimming, but little orientation change. In the second category we scored, cells approached from the sides and first encountered the oral lobes posteriorly. These mostly elicited larger-scale ciliary reversals, often involving the anterior lobes as well as the oral lobes. Consequently these events involved, on average, greater orientation change (Fig. 5A, plot 3).

The most dramatic events followed the arrival of prey across the dorsal surface of the larva (Fig. 1A, Fig. 5D-F and Video 6). In these cases, the prey cell followed a path toward the posterior end of the body, whereupon a reversal of the oral lobes and anteroventral lobe caused the larva to orbit the prey item, in effect swimming the entire larva around the parcel of food-containing water. These events involved on average >90 degree turns, even complete tumbles (Fig. 5A, plot 4; note that many such records could not be measured because the tumble left the focal plane). Such events likely represent most of the tumbles observed while watching these larvae swimming freely in culture bowls.

All three larval morphotypes that we observed to feed followed similar patterns, but varying in degree. The *Pseudoceros* type (Fig. 5B,D,E), the smallest, executed the most dramatic tumbles, whereas the long-nosed type had only subtle turns despite long-duration reversals. Hence our measured direction changes (Fig. 5A) include only the “Winnie-the-Pooh” types, only sequences in side view at 20x, and only sequences with successful capture (fumbles, predictably, often involved extended tumbles). Within this set of 47 recordings, in 34 cases we could score the presence or absence of reversal on oral, anteroventral, and anterodorsal lobes. The greatest average direction change occurred when oral lobes and anteroventral lobes reversed together (102°, n=13). Sequences in which only the oral lobes reversed underwent predictably modest turns (37°, n=7). In cases in which both oral lobes and anterodorsal lobes reversed, regardless of whether the anteroventral lobe reversed, even more modest turns resulted (26°, n=14); of these, the few sequences in which oral and anterodorsal lobes, but *not* anteroventral lobe, reversed had the smallest direction change (14°, n=3).

Restricting these behaviors within range of a single focal plane required confinement by a slide and coverslip, both within a body diameter; hence, these numbers surely underestimate what would transpire when swimming far from a boundary. Indeed, when recording in cuvettes, we observed many misses that seemed to result from an inadequate turn or inability to create enough suction against the boundary drag to draw in the targeted prey. Larvae tethered far from a boundary were not observed to successfully capture any prey – unsurprising, given that they can barely turn at all. We therefore tried to learn, by recording at low magnification in a dish and far from the coverslip, whether the description given herein applies in the absence of a nearby boundary. It is this effort that resulted in sequences like those in Fig. 5B-F (and Video 6). Although the catch per unit effort for the microscopist is low and it is much harder to resolve details of ciliary behavior, these recordings confirm that our description applies qualitatively to freely-swimming larvae. A notable feature is documented by Fig. 5F’: this panel shows a 30-frame (60 ms) average of successive frames in mid-turn, in the laboratory reference frame. Most of the larval body is naturally blurred out – it’s moving relative to the microscope. However the oral lobe appears as if in sharp focus. To appear so, it must have been nearly stationary while the larva rotated around it mid-water. In other words, the oral lobe operates like a caterpillar tread. Moreover, the “teeth” of this tread consist of the metachronal wave of ciliary beat! Thus the reversed ciliary beat in fact operates like a worm drive with the water itself as the spur gear.

Some Müller’s larvae grow to large size with elaborate tentacle arrangements; does the suite of prey capture behaviors remain the same as the body changes? We caught few larger individuals in plankton, all of which appeared to conform to either the *Pseudoceros* or Long-nose morphotypes. These were challenging to observe feeding because of size and bodily complexity, but the successful recordings confirm that ciliary reversals on oral lobes, anterior lobes, or tentacles accompany prey capture and lead to tumbles (Videos 7 and 8). The sole novelty we detected in larger forms is that tentacles can be flexed either at their base or along their length to deflect passing streams toward oral lobes or to re-orient the body.

### 3.5 Digestion and escape

Unconstrained larvae fed avidly on *Rhodomonas* and diffidently on *Prorocentrum*, but mostly ignored *Tetraselmis, Dunaliella* and *Isochrysis*. They were occasionally observed to ingest *Pyramimonas*, and in a few cases caught larger dinoflagellates of the *Protoperidinium* type and even, in one instance, a small ciliate which promptly escaped or was spat out. Two individuals, when found in a plankton sample, contained centric diatom tests. Feeding larvae rapidly acquired the characteristic magenta color from feeding on *Rhodomonas lens* or *abbreviata*. However, we noted that digestion appears incomplete. Actively-feeding larvae regularly regurgitated masses of partially-digested cells, bright green in color. Regurgitation was preceded by a change in body form: the stomach could be seen to constrict in the larval midsection, and expand in the anterodorsal lobe; color was transported to this newly-expanded lobe, along with clear circulating fluid. This appears to represent the partitioning of fluid chyme from indigestible solids. Thereafter, larvae assumed a tight orbit, a bodily shudder, and thence a mass of several to hundreds of cells worth of debris emerged from the mouth (Video 9). No intact cells could be seen when these pellets were examined at high resolution, only bright green blocks roughly the shape of the plastid within the algal cells. It may be that Müller’s larva cannot digest the storage products of cryptophytes, or it may be that they forego to do so when sufficient prey are available. If the former, it may explain why others have found that unusually high prey density is required to sustain Müller’s larvae in lab culture (Allen et al., 2017).

We also observed that cryptophytes frequently attempted to escape. Escape attempts are evident in high-speed recordings as one or more rapid (~5 ms) jumps of 30-40 microns against prevailing currents. Such jumps were observed once prey found themselves between the oral lobes, or approaching the mouth, or even once in the stomach; by inference, either direct contact by many cilia or the chemical signature of imminent digestion prompts these cells to fire their large ejectisomes. The Müller’s larvae we recorded are small compared to many other ciliated invertebrate larvae, and the cleft between oral lobes is little deeper than the distance that can be crossed by cryptophyte’s ejectisome-mediated jumps (compare to previous account for the nemertean pilidium: von Dassow et al., 2013). Many escape attempts were therefore successful, although the larva also sometimes succeeded in recapturing the cells (Fig. 6 and Videos 10, 11, and 12). Our impression, however, is that larvae may have often missed recapturing cells due to fluid drag that limited their motion in coverslipped preps; they may not miss as many marks in more natural circumstances.

**Figure 6.**
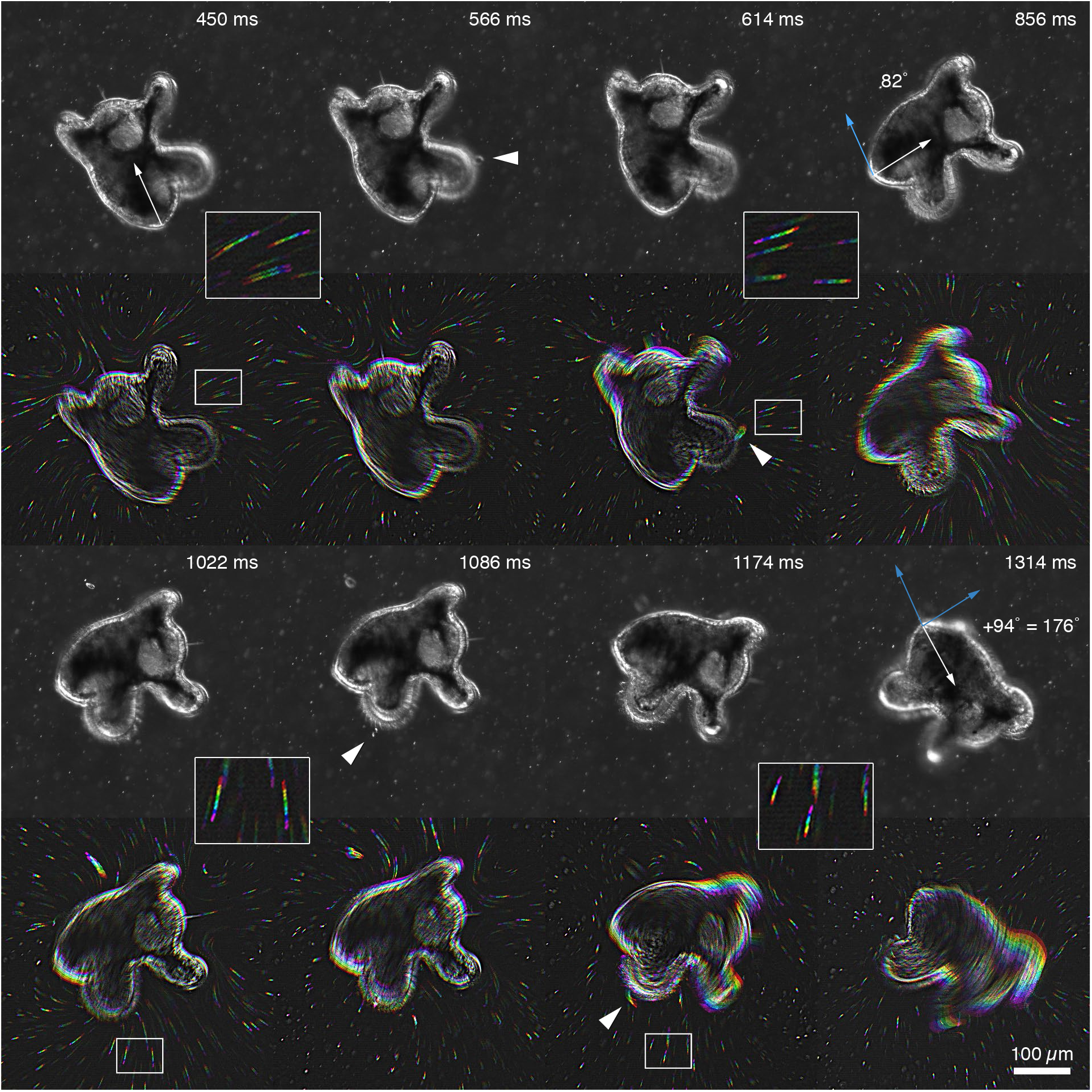
Dealing with prey escape. Frames from a recording of a “free-swimming” larva (>1 body diameter from boundaries) capturing a single *Rhodomonas abbreviata* (white arrowheads), which is initially internalized but then jumps out from between the oral lobes (sixth frame), only to be recaptured with a second reversal and turn. Each raw frame is matched by a high-pass filtered, rainbow-coded version to illustrate flow: the rainbow traces correspond to the 48 ms immediately preceding the unprocessed frame. Insets are 2.5x blow-ups of boxed regions within the draw of the oral lobes.

### 3.6 Bar-coding wild-caught larvae

Whose larvae did we record? The only polyclad encountered frequently in our sampling area that develops via Müller’s larva is *Pseudoceros canadensis*, which feeds on the compound ascidian *Distaplia* and lays eggplates on the underside of the ascidian colony. Indeed, *Pseudoceros* larvae hatched from eggplates conform to the behaviors described here (not shown). Plankton-caught larvae included four clearly-distinct morphotypes (Fig. 7). However, DNA barcoding with 16S and COI showed that of the three morphotypes we used for highspeed recording, only one of 22 barcoded individuals matched *Pseudoceros* (collected from ascidians in the same marina in which larvae were caught). Others did not have a species-level match in Genbank (some individuals failed barcoding), and the most common morphotype was resolved by COI into two clades, distinguishable to us only by a slight color difference. Casual sampling in the marina retrieved only one other polyclad, a species of *Notocomplana* that develops to a crawl-away juvenile, so the parents of these larvae remain as yet unknown, although many candidates are described from this region (Hyman, 1953).

**Figure 7.**
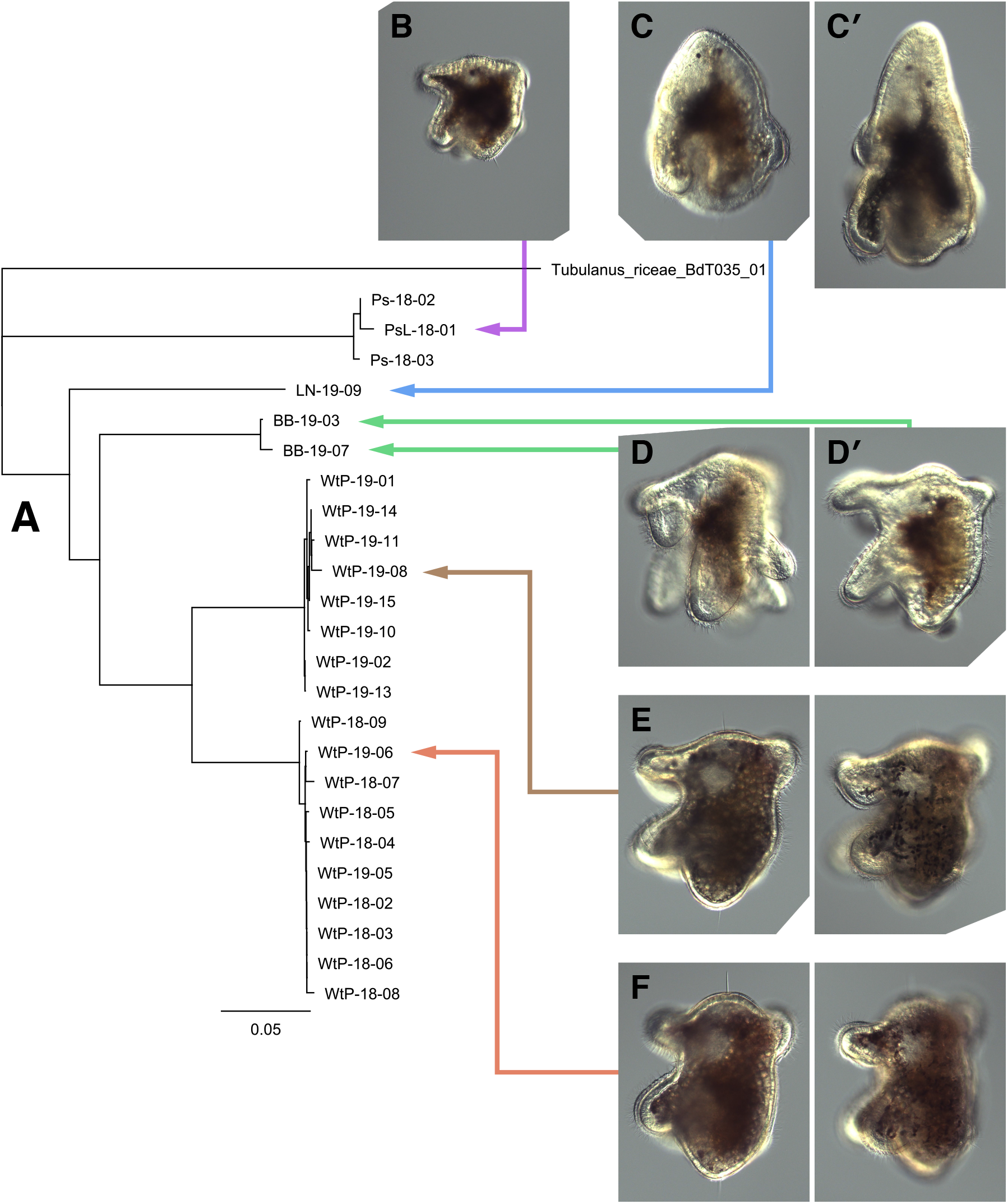
DNA barcoding reveals five possible species. A) Neighbor-joining tree based on COI sequences from 22 individuals across two different years. Individuals are labeled by morphotype and year of collection and correspond to images: Ps=Pseudoceros (B); LN=Long-nose (C and C’, an un-sequenced individual swimming with elongated posture); BB=Broad-bill (D, D’, showing broad and divided anteroventral lobe); and WtP=Winnie-the-Pooh (E and F, the same individuals at medial and superficial focal planes). The most common morphotype in tows is the “Winnie-the-Pooh” variety (E,F), which is here resolved into two sister clades. Only a single specimen (B) yielded sequences that matched locally-collected *Pseudoceros* cf. *canadensis* from *Distaplia;* another singleton is the “Long-nose” type (C,C’; DNA could not be amplified from all larvae of types *Pseudoceros* and Long-nose that we preserved). The Broadbill morphotype (D,D’), was not willing to feed on camera.

### 3.7 Conclusions

Like the fictional clean-up robot Wall•E, Müller’s larva trundles along on treads, spotting and retrieving items it passes, then compressing the aggregate into a discard mass while retaining bits and pieces of potential use (Stanton, 2008). Müller’s larva combines familiar elemental strategies in novel ways to capture unicellular algal prey. Like the nemertean pilidium (von Dassow et al., 2013) and phoronid actinotroch (Strathmann and Bone, 1997), and *unlike* many better-known ciliary feeding larvae), Müller’s larva captures algal prey one by one after detecting individual cells passing near the ciliary bands. Also like the nemertean pilidium and the phoronid actinotroch, individual prey appear to be detected as they meet the ciliated band, possibly by collar cells emdedded therein, prompting muscular responses both near the detection zone and across the body. Unlike the pilidium, though, but reminiscent of echinoderm dipleurulas instead (Strathmann et al., 1972; Strathmann, 2007), transient ciliary reversals redirect otherwise posterior-ward fluid streams. In Müller’s larva, however, these reversals involve heavily multiciliated cells more similar to those in the spiralian prototroch or the cyphonautes’ corona. Moreover the redirection in question is not mouthward, as it is in dipleurulas and other upstream collectors, but actually away from the mouth: anterior lobe reversal nominally pulls water away and out of the oral zone, as does the reversal of the oral lobe’s beat. This works because the oral lobes are normally beating almost toward each other; hence colliding streams drive outward flow during normal swimming. Upon reversal of beat, flow reverses, thus drawing water from afar toward the mouth along a narrow approach fan. Perhaps most worthy of comment is that this strategy depends on the larva rapidly adjusting its body orientation and posture such that detected prey lie within the approach slot when oral lobe cilia reverse. How particles are ultimately internalized is not entirely clear, because it is hard to see, but it appears to involve the activation of a loop of ciliary band posterior to or even entirely surrounding the mouth itself.

A curious consequence of the feeding mechanism used by Müller’s larva is that each prey item necessitates a change in swimming direction. As for chemotactic bacteria or foraging predators that lack sensation at a distance, a turn rate that varies directly with attractant gradient or prize encounters may facilitate “prey-taxis”, i.e., enable swarms of predators to track prey concentrations (Karieva and Odell, 1987). When cultured in dishes with cryptophytes for food, our Müller’s larvae commonly are found in flocks at one edge, or near the bottom where inactive algal cells accumulate. It has not been possible to distinguish whether common phototaxis or other culture artifacts are the basis for such observations, but it seems equally plausible that their feeding behavior could be the cause. It would be of interest to evaluate whether these and other zooplankters that use a one-by-one capture mechanism with a transient flow-field disruption – actinotrochs, pilidia, even dipleurulas – achieve natural colocalization with prey blooms. This seems most likely to work for Müller’s larvae, because they are so compact-bodied in comparison with the balloon-like larvae of blastocoel makers.

## Supplemental figure legends

**Supplemental Figure 1.**
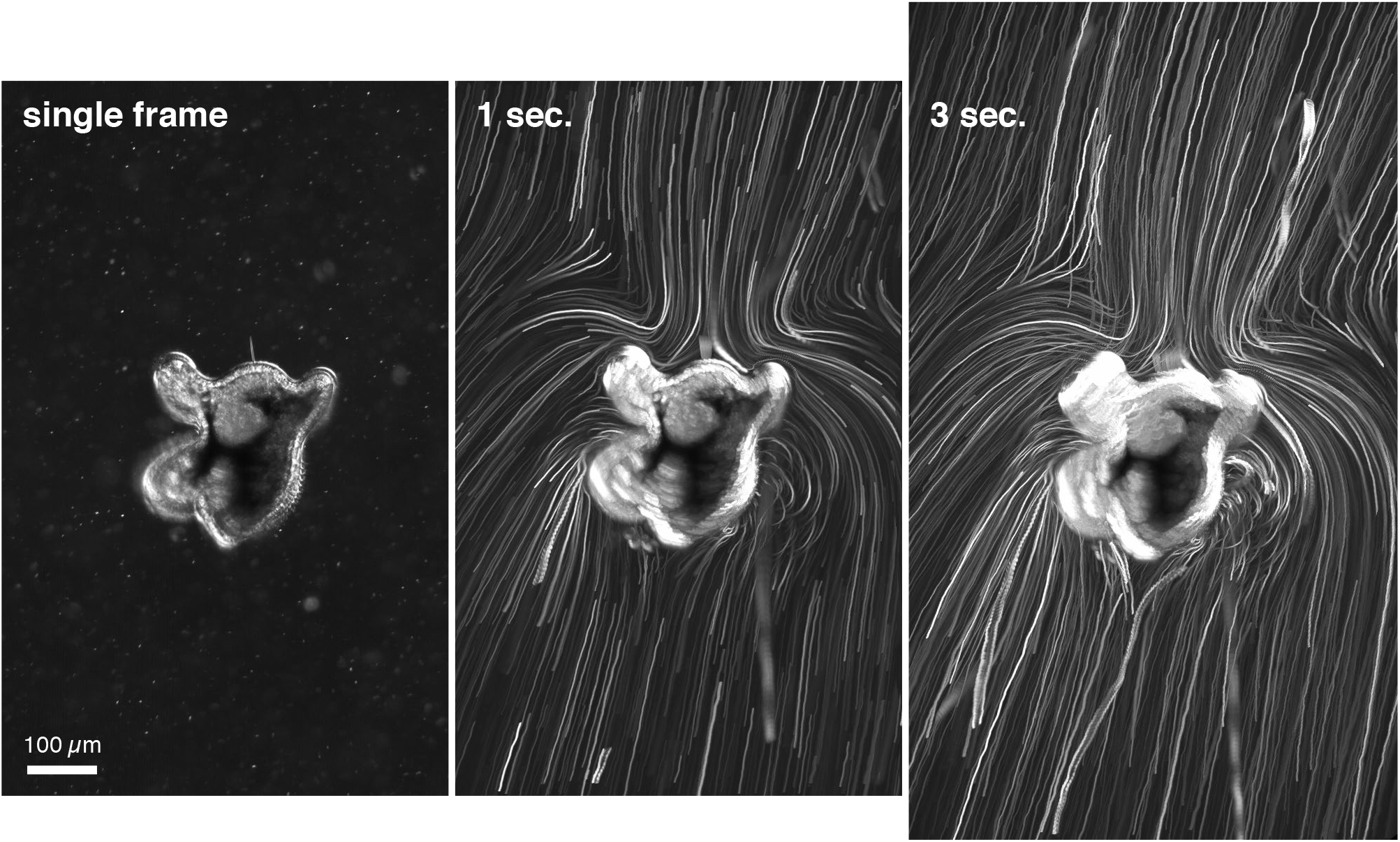
Pathlines around freely-swimming Winnie-the-Pooh type larva in a coverslip-bottom dish, seeded with dilute milk suspension, and further than one body diameter from any boundary. Left panel is a single frame; middle panel is a maximum projection of 500 frames (=1 sec.) after image registration; right panel is a maximum projection of 1500 frames (=3 sec.). Features of note include 1) the slow convergence, then sharp divergence, of pathlines approaching the anterior lobes; 2) vortices along the dorsal surface, where the dorsal ridges of ciliary band drive sharply converging flow; and 3) greater density of pathlines passing either oral lobes or dorsal surface, both of which are apparently reactive, compared to the larval posterior.

**Supplemental Figure 2.**
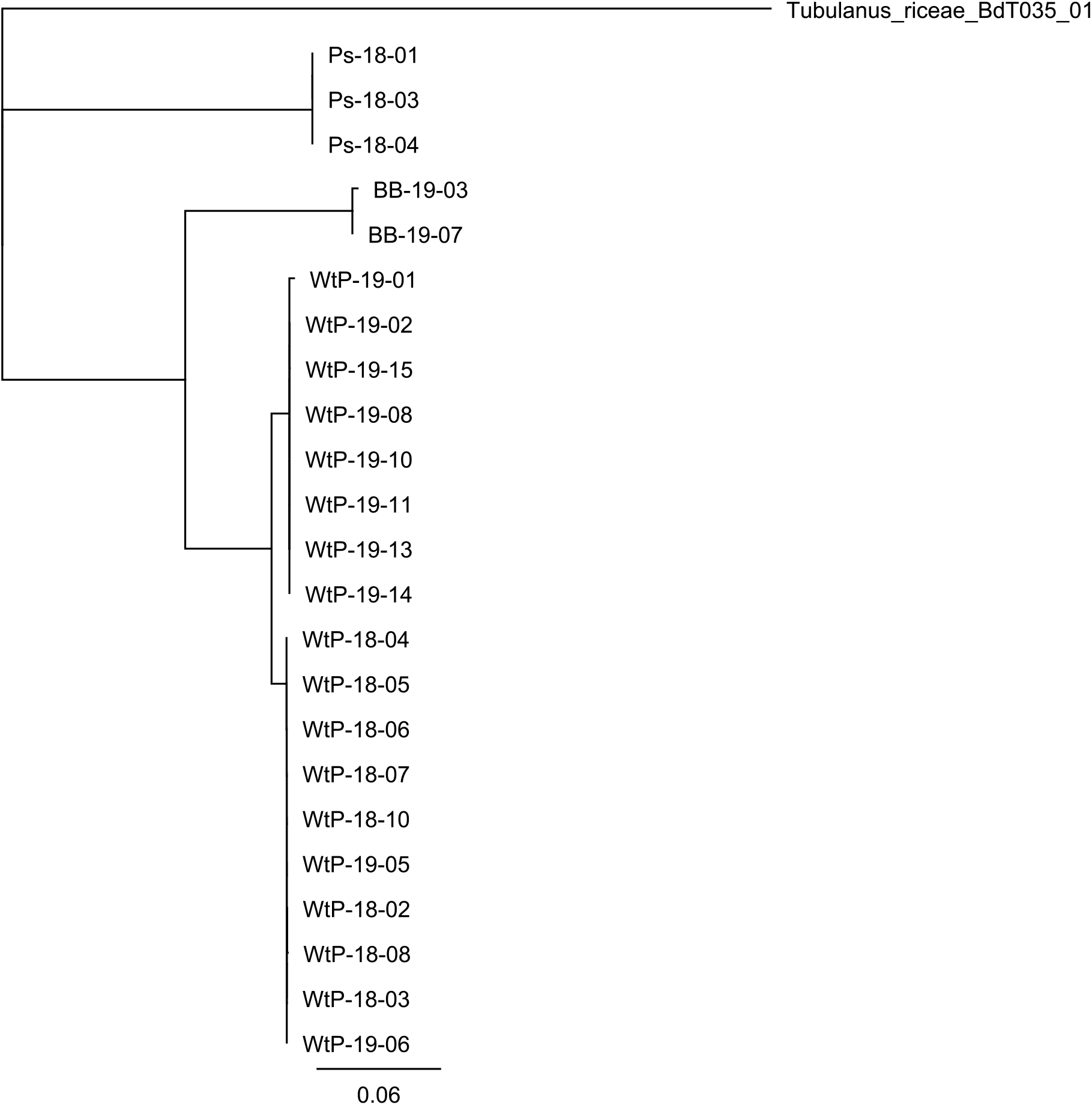
Neighbor-joining tree based on COI sequences from 22 individuals across two different years. Individuals are labeled by morphotype and by year of collection, as in Figure 7.

## Supplemental video legends

Video 1. Standard capture sequence, side view, in a cuvette supplied with *R. abbreviata*. Corresponds to Figure 1A. Prey is encountered dorsally, whereupon detection elicits ciliary reversal in anteroventral and oral lobes. Larva consequently tumbles, ultimately turning completely to move mouth to meet prey item. Video frames registered to approximate larval reference frame.

Video 2. Standard capture sequence, posterior view, in a cuvette supplied with *R. abbreviata*.Corresponds to Figure 1B. Prey approaches along diagonal pathline passing oral lobe. Upon detection both oral lobes reverse ciliary beat and move slightly apart; prey is thus drawn between them instead of continuing ventrally.

Video 3. Standard capture sequence, side view, in a cuvette supplied with *R. abbreviata*. This sequence includes two successful captures within ~1 sec. of real time. Both captures involve reversal of oral and anterior lobes. Note also the onset of ciliary beating in a post-oral loop of the ciliated band, which is otherwise at rest. No corresponding figure.

Video 4. Capture sequence, side view, in a dish with *R. abbreviata* and dilute suspension of milk particles to visualize flow. Corresponds to Figure 4A. Four versions play: the original image, registered to convert to larval reference frame, processed by high-pass FFT, and projected over 60 frames (=120 ms) to demonstrate change in near field flow accompanying ciliary reversal. In this instance, reversal takes place on both anteroventral and oral lobes. Note that this animal is swimming further than a body length from any boundary, as shown from the absence of out-of-focus fixed-position objects in the image.

Video 5. Capture sequence, dorsal view, with *R. abbreviata*. Corresponds to Figure 4B. Focal plane cuts through posterodorsal lobes. Prey encounters ciliary band of left-side lobe (actually right side; mirror image from inverted microscope), transiently deflecting one of the non-beating cilia. This appears to trigger reversal on opposite lobe (and likely elsewhere), but *not* in the zone which encountered prey. Consequently larval body rotates to align prey with approach fan through oral lobes.

Video 6. Characteristic capture sequences illustrating prey routes and associated behaviors, filmed at low magnification, swimming nearly freely in coverslip-bottom dishes. Corresponds to Figure 5B-F. Sequences include both *Pseudoceros* and Winnie-the-Pooh morphotypes.

Video 7. Capture sequence involving large, tentaculate Pseudoceros-type larva swimming in a cuvette with *R. abbreviata*. Prey passes posteriorly along outer side of oral lobe; oral lobes and anteroventral lobe reverse, and oral lobes swing apart as prey is drawn mouthward. No corresponding figure.

Video 8. Five low-magnification capture sequences involving large, tentaculate *Pseudoceros*-type larvae swimming freely in dishes. Segments 1, 3, and 4 feature the same individual; sequences 2 and 5 are a different individual. These larvae were wild-caught at this stage, with eight lobes – two anterior, two oral, and four posterior – as none of the larvae reared from collected egg plates could be raised past six lobes. (No Winnie-the-Pooh type larvae developed beyond six lobes either, at which point they metamorphosed.) In each segment, the apparent primary prey item is highlighted by an arrow. In some cases, other prey are captured simultaneously. Segment 1 features a simple capture in which anteroventral and oral lobes undergo reversal, pivoting the larval body around a single cell. In the other segments, prey items first encounter a tentacle, which flexes while maintaining ordinary beat. In some cases it is clear that other tentacles on the opposite side of the body undergo reversal, presumably to promote bodily rotation. It is remarkable that the prey item, as the larva maneuvers, follows trajectories that take it as much as 100 microns or more from the larval body, and is nevertheless accurately caught. No corresponding figure.

Video 9. Ejection of undigested material. Initial sequence shows normal posture and body shape in a Winnie-the-Pooh type larva that has been feeding avidly on *R. lens*, and hence has a bellyful of magenta. Second and third sequences show posture and body shape as the larva prepares to eject a waste mass: the greatly expanded anterodorsal lobe contains magenta color and seemingly fluid stomach contents, whereas ejectate mass is nearly grass-green. Such masses contain the remains of many hundreds of cells. No corresponding figure.

Video 10. Capture sequence involving attempted escape. Larva is swimming nearly freely in a coverslip-bottom dish with *R. abbreviata* and a dilute milk suspension. Initial capture involves ~90° rotation. Shortly thereafter, the prey jumps several times to exit the mouth, appearing between the oral lobes. Another reversal and concomitant turn accomplishes recapture. Video includes both original images and FFT band-pass filtered, rainbow-coded version to show flow orientation around the larval body. Corresponds to Figure 6.

Video 11. High-magnification sequence involving attempted escape. Larva is swimming in a cuvette with *R. abbreviata*. Initial capture features an especially clear instance of reversal along the oral lobe. Prey promptly jumps out as ciliary beat is restored, but is immediately recaptured as oral lobes reverse beat again. Prey executes another jump sequence as oral lobes continue reversed beat, but is carried within reach of anteroventral lobe, and is again recaptured. No corresponding figure.

Video 12. Successful escape. Larva is swimming in a cuvette with *R. abbreviata*. After capture, prey jumps out but is promptly retrieved. Second escape attempt is successful. No corresponding figure.

---

**Supplemental Table 1.**
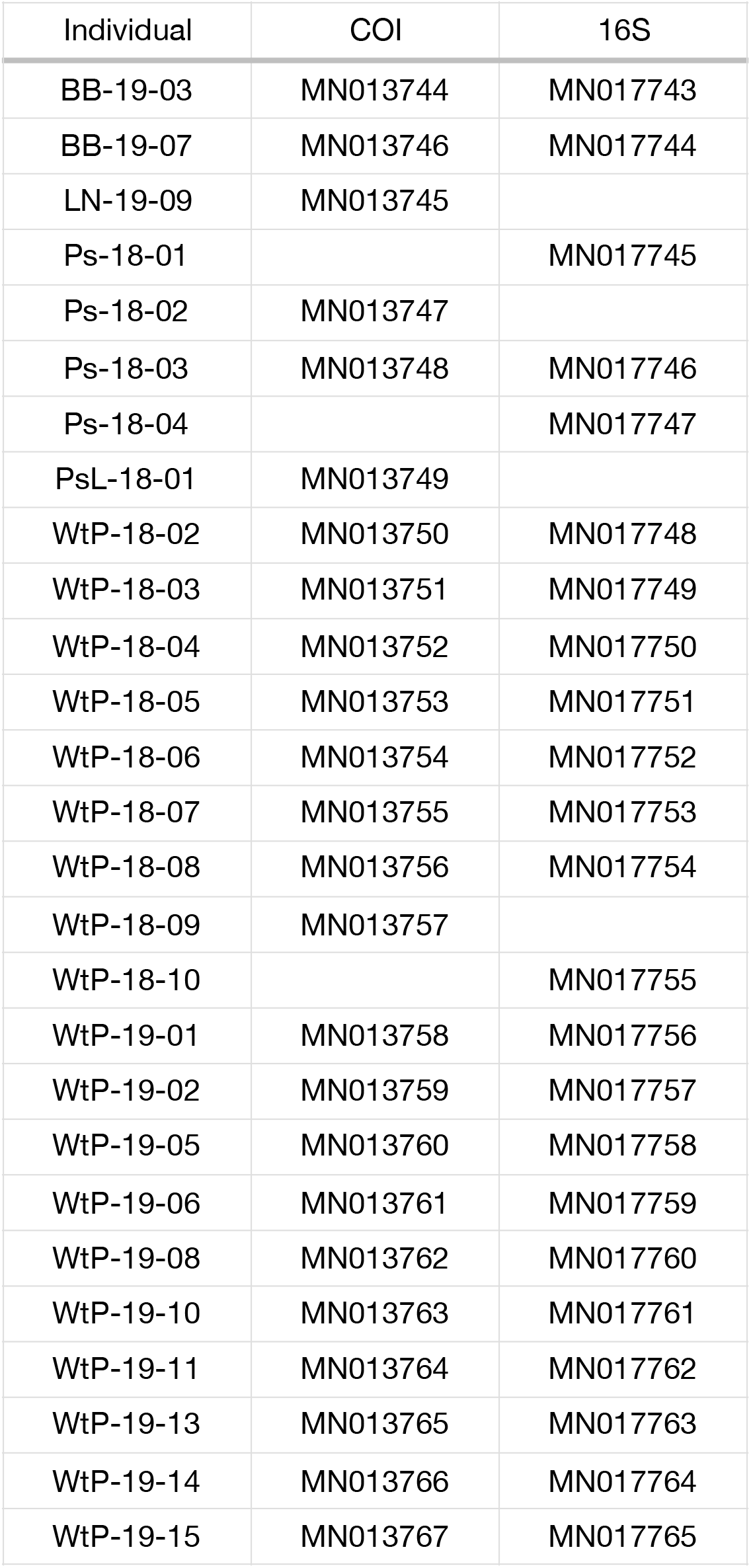
Accession numbers for larvae and adults bar-coded; corresponds to Figure 7.

